# Morph-specific selection pressures drive phenotypic divergence in a color polymorphic bird

**DOI:** 10.1101/2024.10.18.619091

**Authors:** Arianna Passarotto, Moritz D. Lürig, Esa Aaltonen, Patrik Karell

## Abstract

There is a long tradition of using heritable color polymorphisms in natural populations to study selection, gene flow, and other evolutionary processes. However, we still have only limited knowledge on how continuous color variation within genetically discrete morphs affects selective dynamics, which narrows our understanding of how color polymorphisms persist. Our comprehensive analysis of 43 years of plumage color scores from a bi-morphic Finnish population of tawny owls (*Strix aluco*) reveals that intra-morph variation is substantial, but also unexpectedly dynamic. We show that both morphs recently diverged in their plumage coloration: while the brown morph is on a steady trajectory toward more intense plumage pigmentation, the gray morph has recently shifted toward lighter plumage, following a series of extreme winters. Evidence suggests that this divergence is driven by the brown morph’s higher reproductive success and greater plasticity in response to seasonality, while the gray morph, with its more genetically determined plumage color and limited standing variation, has a reduced capacity to respond to selection and track new phenotypic optima. Our study confirms the presence of substantial and dynamic phenotypic variation within genetically discrete color morphs, and demonstrates its relevance for the evolutionary potential of populations to respond to environmental change.

## Introduction

Color variation in animal populations spanning spatially or temporally variable environments is a classic research theme in ecology and evolutionary biology ^1^. Consistent intraspecific variation in color often stems from color polymorphisms, genetically based color variants, whose expression is mostly or completely insensitive to the external environment ^2,3^. Accordingly, color polymorphisms are often (but not always, e.g.^4,5^) determined by Mendelian genetics, with alleles segregated across only a few loci ^6,7^. Based on these assumptions, color polymorphisms have long been used as visual markers of genetic variation, thereby providing a simple but effective tool to study a variety of evolutionary processes in a range of different organisms ^8–10^. However, within what are considered discrete color morphs, there is mounting evidence for the presence of continuous variation in coloration ^5,11–13^, which is thought to be typically controlled by complex arrays of genes (reviewed in ^14^), and moreover can be sensitive to environmental variation - i.e. phenotypically plastic ^15,16^. The presence of intramorph continuous color variation not only complicates their use as visible markers for genetic variation ^10^, but also our understanding of the evolutionary processes that generate and maintain color polymorphisms in natural populations ^5,16,17^.

The persistence of color polymorphisms within populations implies a balance of selective forces that keep the frequencies of each morph relatively stable over time ^3,18^. Mechanisms that lead to equivalent average fitness among color morphs include negative frequency-dependent selection ^19,20^ and variation in selection across environmental gradients (Fig. 1A) ^7,21^. When external factors such as climate change alter these gradients, they also reshape the fitness landscape of polymorphic populations, which can have potentially dramatic consequences for the persistence of the polymorphism ^8,22^: in a fully discrete system, changes to the selective regime would only affect the frequency of the morphs in a population ^17^, whereas in species where the morphs are characterized by continuous, non-overlapping phenotypic differences, within-morph variation may act as fuel for directional selection. Intraspecific variation would therefore enable the morphs to trace new phenotypic optima ^17^ (Fig. 1A), however, only up to the physiological limits - e.g. for color expression set by genetic and developmental constraints in pigment production ^23–25^. Therefore, novel or changing selection pressures might not only affect morph frequencies directly, but also through the distribution of color phenotypes within a morph.

**Figure 1.**
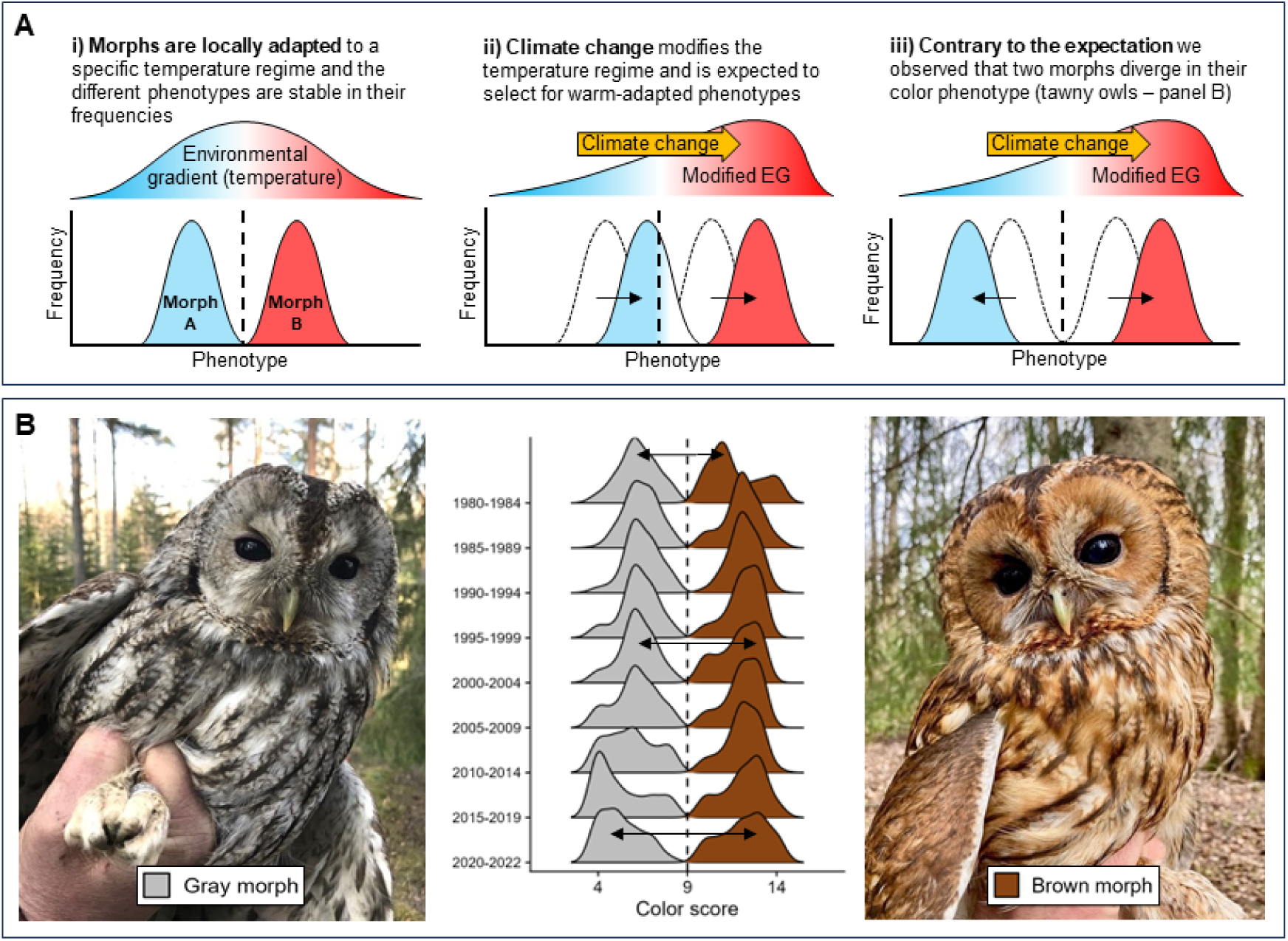
Color polymorphisms under a changing environment. **A:** Schematic illustration of a color polymorphism with continuous variation among morphs adapted to different temperatures: Morph A to cold and Morph B to warm temperatures. Panel (i) shows stable phenotype frequencies around each morph’s optimal color phenotype. The vertical dashed line indicates the physiological limit for color expression due to genetic and developmental constraints. As climate change induces a shift towards higher temperatures (ii), selection favors warmer-tolerant phenotypes. Despite this, Morph A cannot surpass its physiological limit to the production of coloration, leading to a decline in its frequency under warmer conditions. Conversely, Morph B not only increases in frequency but also exhibits a higher occurrence of warm-tolerant color phenotypes within the morph. In the present study (iii) we observed a pattern that does not match this expectation: instead of moving in the same direction along an axis of coloration, the two morphs of tawny owls (*Strix aluco*) diverge in their coloration. **B:** Density plot of plumage color scores from 4 (pale gray) to 14 (dark brown) recorded in the Finnish population between 1978 and 2022. The vertical dashed line marks the cutoff used to assign the two morphs in the population over 40 years (i.e. the putative physiological limit of color expression). While initially both morphs appeared to move toward a more brown-dominated coloration (higher color scores), since the early 2000s the morphs have started to diverge. Photos by Chiara Morosinotto (gray morph) and Gian Luigi Bucciolini (brown morph).

In a Finnish population of color polymorphic tawny owls (*Strix aluco*; Fig. 1B), plumage coloration is highly heritable and expressed in a gray and brown morph following a one-locus/two-allele Mendelian inheritance system with brown dominance ^26,27^. From this discrete genetic basis, both morphs express considerable within-morph variation in plumage coloration (Fig. 1B), through which they might respond to environmental conditions. Specifically, a gray plumage is associated with higher cold tolerance and crypsis in snowy conditions, while brown plumage coloration is expected to be more adaptive to warm winters with little snow ^28^. Previous research indicated a rapid shift in morph frequency from gray-dominated to roughly equal frequencies of both morphs ^26^, which is attributed to relaxed selection against the brown morph in milder snow-poor winters ^26^, and supported by recent work that identified selection on camouflage coloration and plumage characteristics as possible mechanisms ^29–31^. Therefore, it is conceivable that climate change may impose unidirectional selection pressures for plumage phenotypes that are beneficial under increasingly mild winter conditions (i.e. less gray, more brown coloration; Fig. 1A). However, the lack of quantitative studies in wild populations leaves many questions unanswered, for instance, how physiological limits imposed by genetic constraints or low standing variation may affect selective dynamics on color phenotypes.

Here, using a 43-year dataset of plumage color scores, we investigate the extent and possible role of intramorph phenotypic variation for evolutionary change in a long-lived polymorphic bird of prey. Specifically, we test i) for the presence and dynamics of intramorph variation in plumage coloration, ii) whether variation in plumage coloration affects breeding behavior across a range of winter conditions, and iii) whether developmental effects contribute to the observed divergence in plumage coloration. Overall, our study reveals substantial intramorph variation in plumage coloration, whose dynamics are shaped by a complex interplay of morph-specific selection on life-history traits in breeding adults, and differential sensitivities to genetic and environmental factors in juvenile owls that are highly morph-specific.

## Results

### Variation in intramorph plumage coloration

Using standardized and repeatable ordinal color scores (Fig. S1; see Methods for details) from all usable breeding adult encounters (*n* = 1972; Table S1), we detected a divergence in plumage coloration between the two morphs (Fig. 2A), with decreasing color scores in the gray morph (i.e. more “grayness”) and increasing color scores in the brown morph (i.e. more “brownness”). However, while color change in the brown morph was best described by a steadily increasing linear trend in a Generalized Additive Model (GAM 1; Table 1), the pattern in the gray morph was characterized by a hump-shaped form with increasing values during the 1980s, slowly decreasing values between 1990 and 2010, and rapidly decreasing values since 2010, following the occurrence of extremely harsh winters (red annotations in Fig. 2E-H). These morph-specific differences persisted after controlling for observer-specific effects on color scoring, the contribution of immigrating individuals, and comparison between new demographic cohorts (i.e. only first-time breeders new to the population) and the entire (cumulative) population (GAMs S1-2; Table S2; Fig. S2A-C). To confirm the heritable nature of coloration ^26^, we conduct a parent-offspring regression using using linear model with 170 pairs, showing that color score (i.e. quantitative color expression) in tawny owls is highly heritable (H^2^ = 0.83, R² = 0.41; Fig. 3A).

**Figure 2.**
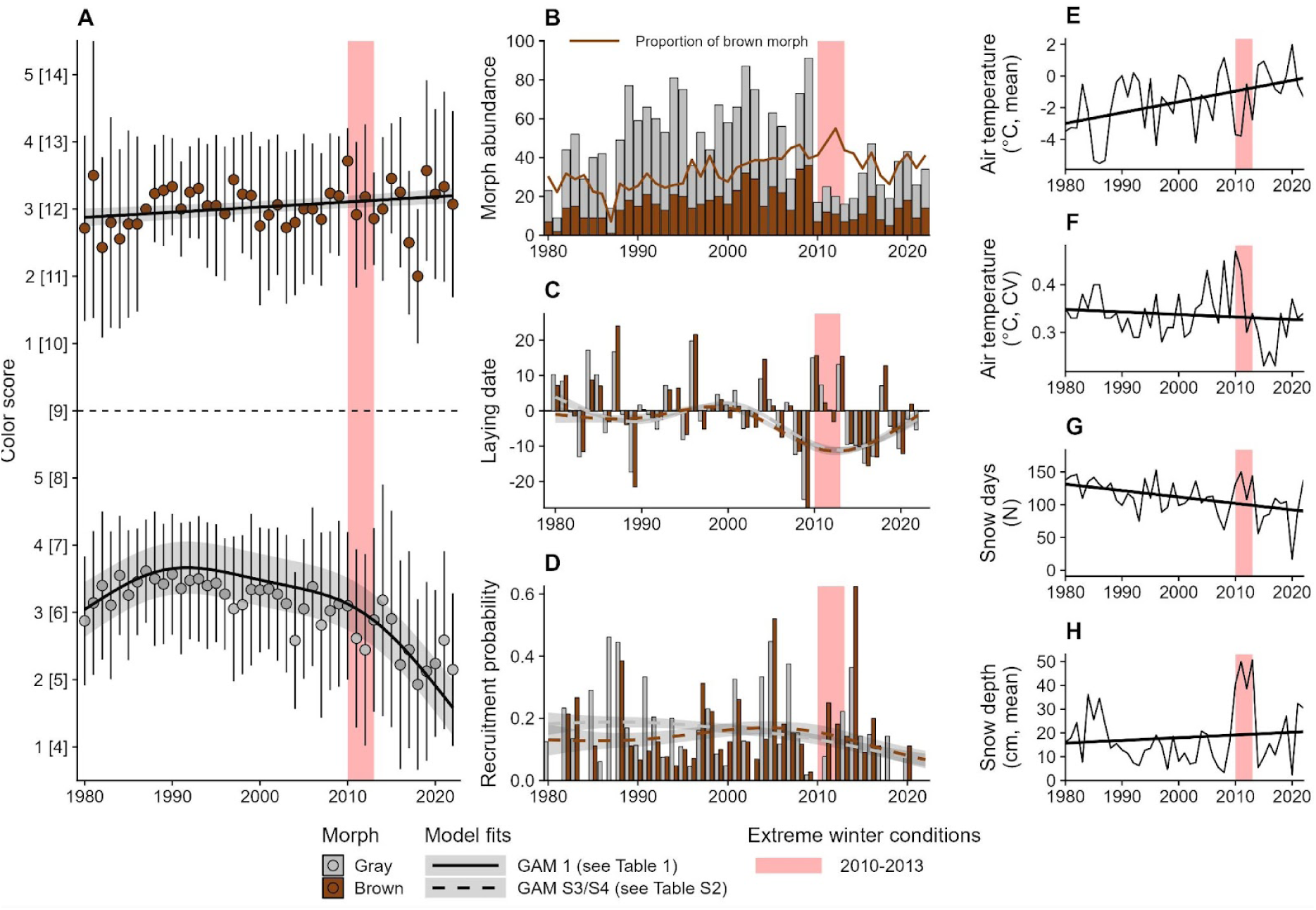
Tawny owl study population between 1980 and 2022. **A:** Color scores of captured breeding adults are highly divergent over the years. Points denote the yearly average of morph-specific color scores using all breeding individuals per each year (mean ± SD, *n* = 1972). Model fit (solid line) stems from GAM 1; shading denotes ± SE of fit. The red annotation depicts a series of severe winters (panels E-F), which resulted in a dramatic reduction of captures (panel B). On the x-axis both relative and absolute (in square brackets) color scores are provided. **B:** Abundances of all individuals per morph (bars) and morph frequencies/proportion of the brown morph over the years (brown line). **C:** Average egg laying date for each morph in the study population, relative to median laying date March 31st (= 0). Dashed lines are morph specific model fits from a Generalized Additive Model (GAM) that found no differences between the morphs in average laying date over time (GAM S3; Table S2). **D:** Probability of recruit production per morph (dashed lines) over the years. Dashed lines are morph specific model fits from a GAM that found a reduction in recruitment success in the gray morph, but not the brown morph (GAM S4; Table S2). **E-H:** Information on winter conditions expressed as the mean air winter temperature, coefficient of variation (CV) of air temperature, the sum of snow days, and mean snow depth that were collected by the Finnish Meteorological Institute (FMI) at the weather station near Helsinki-Vantaa airport in proximity to the study area. The solid black lines are the non-standardized model estimates from a path analysis (SEM 1; Table S3).

**Figure 3.**
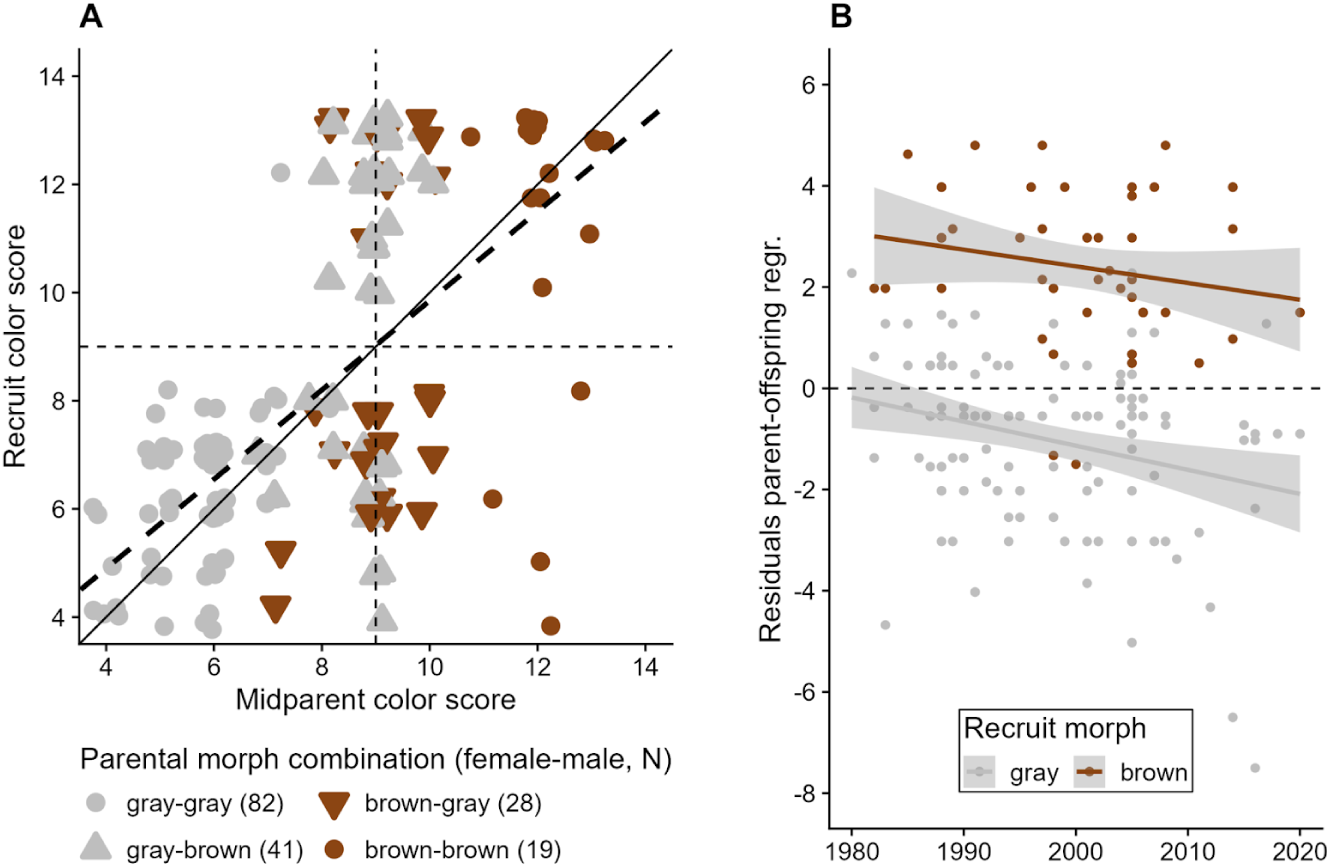
Heritability of plumage coloration in tawny owls. **A:** Parent-offspring regression (*n* = 170), using the midparent values for absolute color scores. The thin dashed lines denote transitions between morphs of parents of offspring, where color scores below 9 indicate gray morph, while higher color scores represent brown morph. Symbols depict different parental morph combinations. The thick dashed line denotes the regression line (coefficient = 0.83, R² = 0.41, *P* < 0.001); the solid line is the 1-1 line, for comparison. Data show the presence of brown recruits originated from gray-gray parents despite under a Mendelian inheritance the gray morph should be homozygous recessive. This could be due to extra pair paternity, which is present in this species although uncommon. **B:** Residuals of the parent-offspring regression over time, grouped by recruit morph. Gray shading shows corresponding standard error.

**Table 1.** Summary table for Generalized Additive Models (GAMs). GAM 1 used absolute color scores (4-14), GAM 2 used relative scores (1-5). Est. df is estimated degrees of freedom, where 1 indicates that a single degree of freedom is required to model the relationship indicating that it is a linear function. Higher values indicate that more degrees of freedom are required to model the relationship, indicating that it is non-linear - e.g., in temporal change of color scores in the gray morph. Significant (0.05 > p ≥ 0.01, *) and highly significant (0.01 > p, ***) results are in bold.

| Model | Response variable | Component | Term |  | Est. df | Ref. df | Chi.sq / F-value | p-value |
| --- | --- | --- | --- | --- | --- | --- | --- | --- |
| GAM 1 | Color score | Linear component |  | morph |  | 1 | 0.352 | 0.553 |
|  |  | Smooth terms | Year | gray | 3.757 | 3.965 | 33.196 | < 0.001 *** |
|  |  |  |  | brown | 1.001 | 1.002 | 3.387 | 0.066 |
|  |  |  | Observer | gray | 1.958 | 2.000 | 83.979 | < 0.001 *** |
|  |  |  |  | brown | 1.348 | 2.000 | 2.528 | 0.025 * |
| GAM 2 | Color score | Linear component |  | morph |  | 1 | 0.210 | 0.646 |
|  |  | Smooth terms | Air temp. (Mean) | gray | 1.001 | 1.001 | 6.713 | 0.010 * |
|  |  |  |  | brown | 1.000 | 1.001 | 0.073 | 0.788 |
|  |  |  | Snow days (N) | gray | 3.344 | 3.558 | 6.036 | 0.001 *** |
|  |  |  |  | brown | 1.000 | 1.001 | 0.814 | 0.367 |
|  |  |  | Air temp. x Snow days | gray | 7.753 | 8.905 | 3.439 | < 0.001 *** |
|  |  |  |  | brown | 1.002 | 1.003 | 0.725 | 0.395 |
| GAM 3 | Laying date | Linear component |  | morph |  | 1 | 2.839 | 0.092 . |
|  |  | Smooth terms | Air temp. (Mean) | gray | 3.340 | 3.752 | 47.265 | < 0.001 *** |
|  |  |  |  | brown | 2.254 | 2.771 | 32.961 | < 0.001 *** |
|  |  |  | Color Score | gray | 1.001 | 1.002 | 0.484 | 0.487 |
|  |  |  |  | brown | 1.003 | 1.005 | 0.009 | 0.940 |
|  |  |  | Air temp. x Color Score | gray | 1.739 | 2.188 | 2.295 | 0.086 . |
|  |  |  |  | brown | 1.979 | 2.461 | 1.096 | 0.303 |
| GAM 4 | Recruitment success | Linear component |  | morph |  | 1 | 0.906 | 0.341 |
|  |  | Smooth terms | Laying date | gray | 1.294 | 1.528 | 0.541 | 0.764 |
|  |  |  |  | brown | 1.817 | 2.265 | 3.417 | 0.198 |
|  |  |  | Color Score | gray | 2.645 | 3.191 | 11.170 | 0.013 * |
|  |  |  |  | brown | 2.058 | 2.565 | 2.671 | 0.292 |
|  |  |  | Laying date x Color Score | gray | 1.001 | 1.002 | 2.341 | 0.126 |
|  |  |  |  | brown | 1.000 | 1.001 | 5.835 | 0.016 * |
| GAM 5 | Recruit color score | Linear component |  | morph |  | 1 | 1.665 | 0.199 |
|  |  | Smooth terms | Paternal color score | gray | 2.495 | 3.012 | 3.882 | 0.010 * |
|  |  |  |  | brown | 1.000 | 1.000 | 0.023 | 0.879 |
|  |  |  | Maternal color score | gray | 1.301 | 1.504 | 6.425 | 0.028 * |
|  |  |  |  | brown | 1.001 | 1.002 | 1.451 | 0.230 |
|  |  |  | Paternal color score x maternal color score | gray | 3.646 | 11.000 | 0.889 | 0.011 * |
|  |  |  |  | brown | 0.000 | 3.000 | 0.000 | 0.726 |
| GAM 6 | Recruit color score | Linear component |  | morph |  | 1 | 0.210 | 0.646 |
|  |  | Smooth terms | Air temp. breeding (Mean) | gray | 1.000 | 1.000 | 1.680 | 0.196 |
|  |  |  |  | brown | 1.000 | 1.000 | 4.431 | 0.037 * |
|  |  |  | Air temp. breeding (CV) | gray | 1.039 | 1.076 | 2.165 | 0.130 |
|  |  |  |  | brown | 1.947 | 2.394 | 1.700 | 0.154 |
|  |  |  | Air temp. Mean x Air temp. CV | gray | 1.000 | 1.001 | 0.064 | 0.802 |
|  |  |  |  | brown | 1.000 | 1.000 | 5.292 | 0.022 * |

### Effects of plumage coloration and breeding phenology on adult fitness

To assess whether and how morph-specific coloration divergence correlates with breeding behavior, we examined color scores in relation to laying date (March 31 = 0) and recruitment success (i.e. the proportion of new breeders originated from the monitored population; immigrants were therefore excluded) as a proxy for fitness. Across both morphs, the owls in the study population have responded to increasingly warm winters with little snow by advancing their average laying date by 10 days over the observed period (Fig. 2C; GAM S3; Table S2). At the same time, the probability of successful recruitment (i.e. new first-breeding adults in the population) has decreased from approximately 20% between 1980 and 2010 to 10% between 2010 and 2020 (Fig. 2D). However, recruitment success in the brown morph did not decrease as strongly as in the gray morph (GAM S4; Table S2), suggesting that brown individuals were able to produce more new breeders over time than gray ones. To investigate possible relationships between changing winter conditions, advancing laying date, and differential recruitment success in relation to color score, we conducted a path analysis using Structural Equation Modeling (SEM 1). We found weak effects of snow depth on color score (more “grayness” with more snow) but no direct effect of color score on either laying date or recruitment success (Fig. 4A; Table S3). The path analysis also revealed that temperature had the highest impact on laying date (approx. 2 days earlier for every degree of warming), while the number of snow days and snow depth had only small effects. However, laying date was not associated with variation in recruitment success, which, moreover, slightly decreased in snow rich winters (Fig. 4A; Table S3).

**Figure 4.**
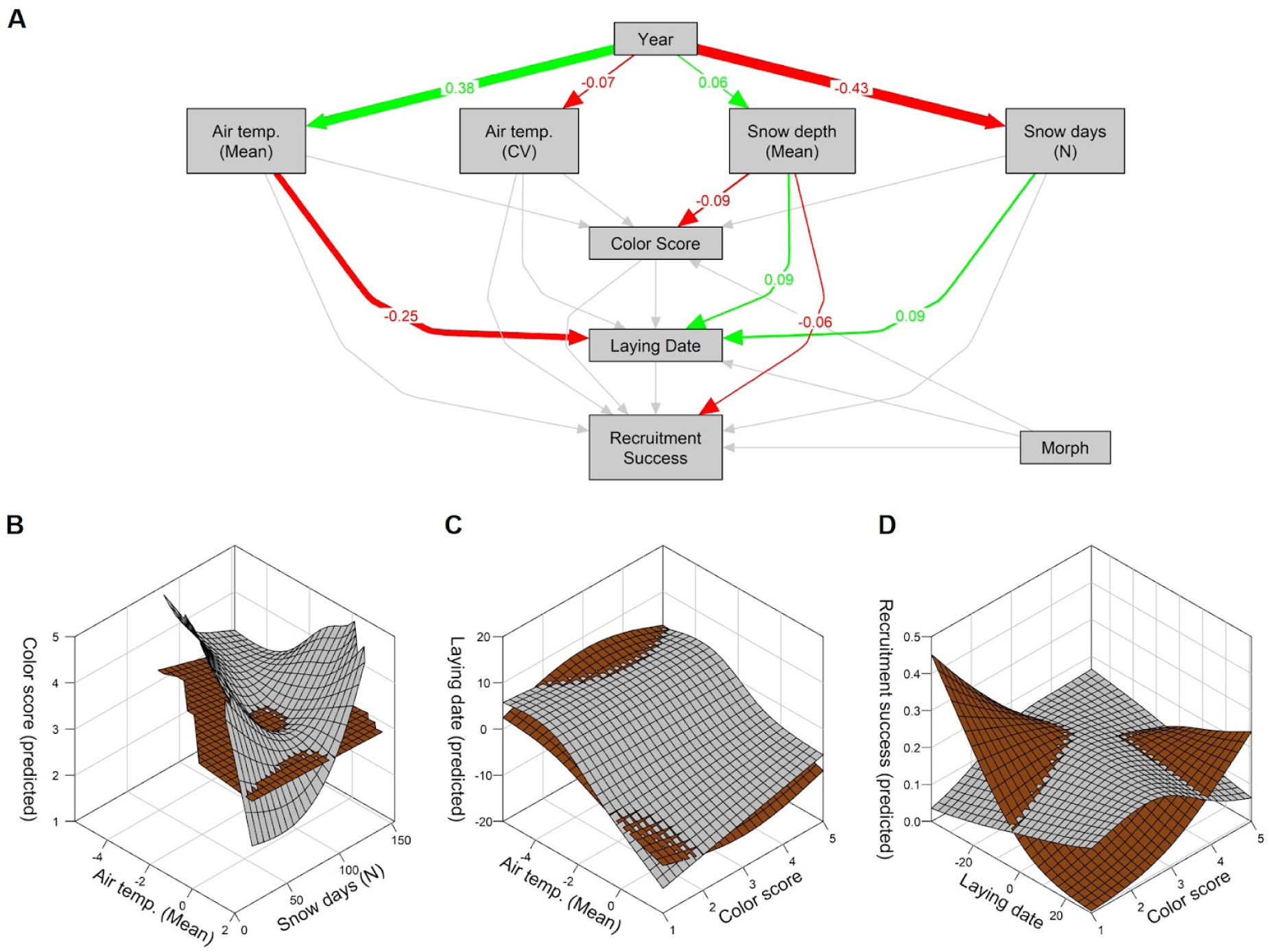
Selection on plumage coloration in breeding adults. **A:** Path analysis based on Structural Equation Modeling (SEM) that includes all observations of breeding adults with known laying date (*n* = 1972, 92.5%), indicating a strong effects of a changing environment on laying date. Colored arrows indicate significant positive (green) or negative (red) effects; gray arrows are not significant effects. The number and width of an arrow show the effect size. Model summary is in Table S3. **B:** Test for interactive effects of morph, snow days and mean winter temperature on the color score of breeding adults using a Generalized Additive Model (GAM 2; Table 1). The model shows that gray individuals were more susceptible to changes in winter conditions. **C:** Test for interactive effects of morph, mean winter temperature and color score on laying date (GAM 3, Table 1) revealed no differences between morphs in breeding behavior according to color score. **D:** Test for interactive effects of morph, color score and laying date on recruitment probability (GAM 4, Table 1). In the brown morph, a close match between breeding adult color scores and laying date increased the likelihood of their offspring recruiting back into the population. In the gray morph, there was a main effect of color score, but no interaction with laying date.

Since SEM allows only for tests of linear relationships without interactions, we used a set of three GAMs (GAM 2-4; Table 1; Fig. 4B-C) to test for possible interactive effects between color phenotype, breeding behavior and fitness. In GAM 2, we used an interactive term with temperature and snow days to assess whether, at the population level and across the entire observed period, plumage coloration was more responsive to environmental changes in one morph over the other. Indeed, the model revealed that the cumulative response to variation in winter conditions was stronger in the gray morph than in the brown one (Fig. 4B). Specifically, gray individuals with higher color scores (i.e. less gray dominated plumage) were breeding in years with either cold winters with few snow days or warmer winters with many snow days, whereas more gray individuals (i.e. low color scores) tended to be breeding in harsh winters with both low temperatures and many snow days. In GAM 3, we used an interactive term with temperature and color score to test whether an individual’s color phenotype would affect the decision when to breed in relation to environmental conditions. We found that, for both morphs, laying date was mostly determined by winter temperature and not by plumage coloration, suggesting that the decision for when to breed is formed independently of the color phenotype (Fig. 4C). In GAM 4, we tested whether the interaction between color score and laying date could predict recruitment success, testing the central hypothesis that light plumage coloration has fitness benefits at earlier laying dates, while darker plumage coloration is expected to be more beneficial when breeding later in the season ^28^. We found support for this hypothesis in the brown morph, where recruitment success was higher when the color score matched the laying date, i.e. when the individual’s coloration was light at earlier laying dates and more intense at later laying dates (Fig. 4D). In contrast, in the gray morph, there was no such interaction and the probability of producing new breeders was only determined by color score, independent of laying date.

### Genetic vs. developmental effects on plumage coloration in recruits

To assess possible temporal shifts in phenotype between parents and offspring, we used a generalized least squares regression model with an autocorrelation term to test for temporal dynamics in the residuals of the parent-offspring relationship, i.e. temporal stability of heritability. We found a small but significant temporal effect of smaller residuals across both morphs (coefficient = -0.23, *P* = 0.002), but no morph specific difference in this effect (*P* = 0.25), which suggests a decrease in similarity between parents and recruits over 40 years in both morphs (Fig. 3B).

Leveraging the same pedigree (*n* = 170), we used a second path analysis (SEM 2) to study the interplay of genetic factors (parental morphs and color scores) as well as environmental effects during the breeding period (same four environmental parameters as in SEM 1) in determining recruit color score. Further supporting the separate parent-offspring regression, SEM 2 (Table S3) showed that plumage coloration in tawny owls is a highly heritable trait (Fig. 5A). Additionally, we found a higher contribution from the maternal than from the paternal side, hinting at the possibility of a sex-linked genetic mechanism. The path analysis also revealed moderate effects of paternal color score and maternal morph on laying date (Fig. 5A), which, however, might be due to the lower sample size compared to all breeding adults in SEM 1.

**Figure 5.**
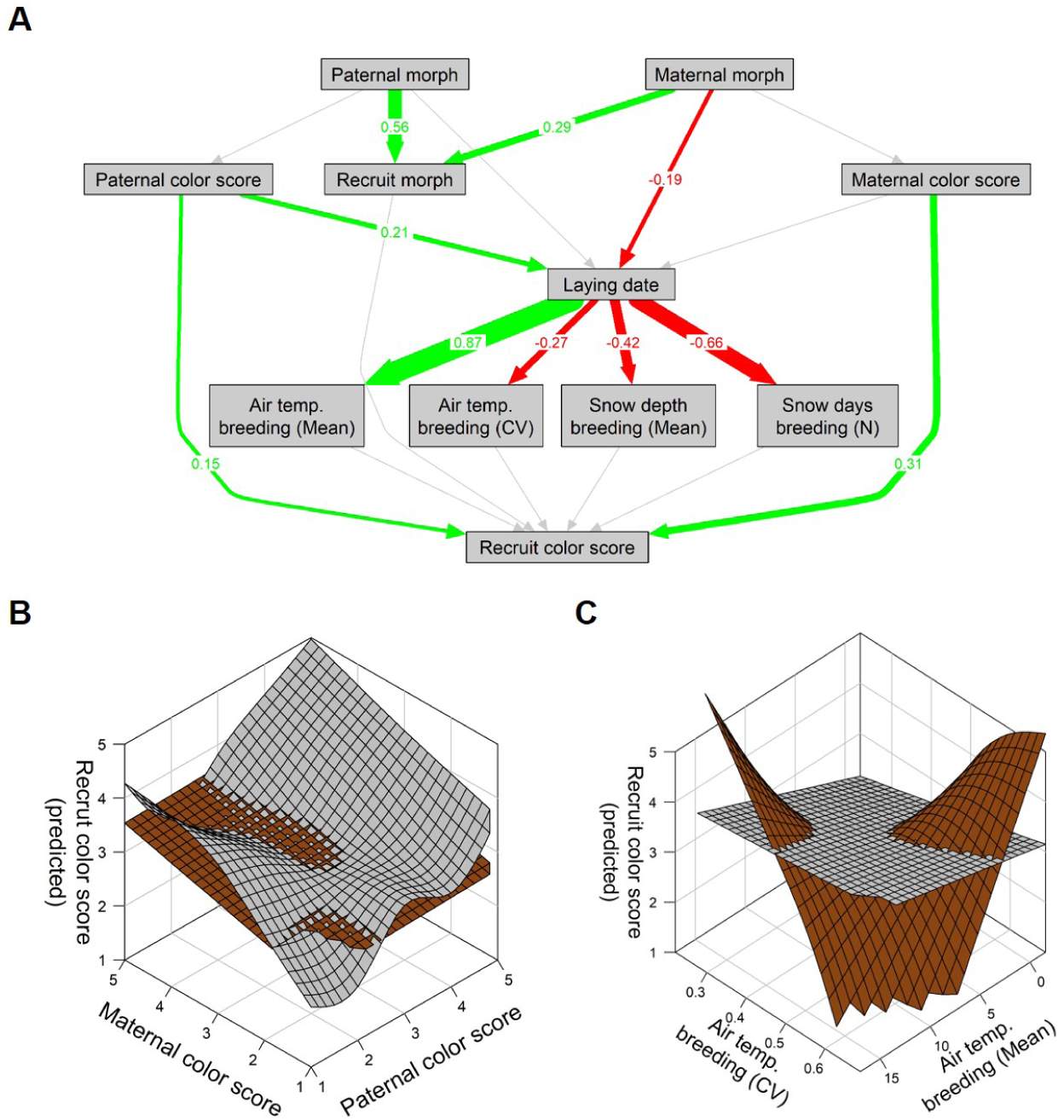
Genetic and development effects on recruit color score. **A:** Path analysis testing for genetic (i.e. paternal and maternal morph and color score) and environmental factors (i.e. laying date, snow depth and temperature before fledging) on recruit color score (*n* = 170), using a Structural Equation Model (SEM 2; Table S2). The path network should be interpreted in the same fashion as the one shown in Fig. 4. Overall, environmental conditions during the breeding period were strongly associated with the timing of the laying date itself. Specifically, a later laying date was associated with higher temperature and both lower temperature coefficient of variation (i.e. CV) and likelihood for days with snow. The most dominant effect on recruit color score is maternal color score. **B:** Model fits from GAM 5 that uses paternal and maternal color score as predictors of recruit color score (Table 1). Parental color scores only affect recruit color scores in the gray, but not in the brown morph. **C:** Recruit color score model fits from GAM 6 that uses mean and coefficient of variation of temperature between laying date and fledging as predictors (Table 1). The model suggests that brown morph recruits are more strongly affected by environmental variation than the gray morph.

We used two GAMs (GAM 5-6; Table 1; Fig. 5A-C) to test for interactive effects on recruit color score. GAM 5 included both parental color scores and their interactive effect, and confirmed a somewhat higher maternal contribution to recruit color score, which is reflected in a higher F-value for the maternal color score (Table 1). A significant interactive effect of parental coloration on recruit color scores in gray but not brown recruits suggests that parental determination of coloration is higher in the gray morph than in the brown morph, which may thus experience a higher degree of developmental plasticity (Fig. 5B). Further supporting this, GAM 6, which tested for effects of environmental variation during early development on color score, revealed a significant interactive effect of mean temperature and temperature variation (i.e. temperature CV) during the breeding period, i.e. between laying date and fledging in the brown morph, but not in the gray morph (Fig. 5C; breeding period = 56 days following laying date [average for the study population]; see Methods for details).

## Discussion

Our analysis of 43 years of field observations indicate substantial within-morph variation in plumage coloration, which, contrary to our expectation (Fig. 1), was highly divergent among both morphs (Fig. 2). On the one hand, following the general expectation of increased pigmentation at higher temperatures (i.e. Gloger’s rule) ^32–34^, we found increasingly intense pheomelanic coloration in the brown morph. On the other hand, and deviating from our expectations, the gray morph followed an opposite trend towards lower color scores, i.e. weaker plumage pigmentation. Our findings suggest that these contrasting changes in a highly heritable trait are likely driven by a combination of morph-specific tradeoffs between life-history strategies and plumage phenotypes ^27^, differential hereditary dynamics, and, putatively, reduced adaptive capacity in the gray morph.

Rapid phenotypic divergence within populations typically occurs when intermediate phenotypes are selected against in the prevailing environment. In tawny owls, warmer winters appear to favor the melanistic brown morph over the gray ^26^, but it remains unclear what other life history traits could contribute to an increase in frequency of the brown morph under warming winter conditions. Brown parents produce fledglings that are in better condition than those of gray parents ^27^. We therefore examined how variation in recruitment success might be linked to differences in plumage coloration, exploring this as a potential driver of the observed divergence within the population. We found that while the decision for when to breed was largely decoupled from coloration of breeding adults of both morphs (Fig. 4C), the probability of successful recruitment was not (Fig. 4D): in the brown morph, recruitment into the breeding population was predicted by a close match between its laying date and plumage coloration, which, following warmer winters, would select for increasingly pigmented recruits. In the gray morph, however, there was no significant interactive effect of color score and laying date, but only a main effect suggesting that intermediate color scores maximize recruitment success across laying dates. This lack of correspondence between pigmentation, environmental conditions at breeding and increase in fitness might be linked to differences in life-history and indicate that plumage color variation in the gray morph is not under the same selective dynamics as in the brown morph. For instance, the gray morph has a longer lifespan and produces more offspring during its lifetime ^35^. Additionally, in other populations, the brown morph was found to have a more constant breeding effort as opposed to a flexible effort of the gray morph, which refrained from breeding more often under unfavorable climate. Therefore, gray adults may achieve higher fitness through alternative strategies and selection may favor plumage colorations that entail higher adult survival, provided that intermediate color phenotypes may have a higher chance of coping with different environmental conditions than either end of the phenotypic spectrum. Although this finding might not fully explain the divergent pattern, we found that in immigrants color scores also tended to decrease over time but not as strong as in the resident breeding pool (Fig. S2B). This indicates that gene flow might counteract the local effects of selection we observed in our population, a possibility warranting further research.

The genetic composition of a population can also be altered by stochastic events that reduce the phenotypic variation natural selection can act upon. Between 2010 and 2013, southern Finland experienced very harsh winters characterized by low temperatures, many snow days, and high snow depth. During these years, the size of the breeding population dropped by about 80%, suggesting increased mortality and a sharp reduction in heritable phenotypic variation. Such bottlenecks may occur without changing morph frequencies directly ^36^, which was also not the case in our population (Fig. 2B), but indirectly by reducing the available heritable phenotypic variation, and thus the potential for evolutionary change ^37^. Accordingly, severe winters may have selected against more pigmented gray individuals, which, along with the high heritability of plumage coloration, might explain the accelerated reduction in average color score after 2013. Therefore, the gray morph may now lack sufficient variation to respond to selection for higher production of pheomelanin under a changing climate ^26^. This pattern is supported by a recent genomic study that identified the genetic basis of cold tolerance in the gray morph and found temporal shifts in the allelic frequencies of the candidate loci’s genotype towards an excess of homozygotes, i.e. individuals of the gray morph, hinting at the possibility of stronger selection among gray owls ^28^. The distribution of plumage coloration in the brown morph, however, did not appear to be affected by the series of extreme winters. Therefore, the brown morph may be able to compensate for phases of low variation through higher phenotypic plasticity (Figs. 3 and 5), making it more resilient when facing climate change and, in general, increasing directional environmental change. This might be also partially due to different dispersal behaviors, which can affect the gene flow by replenishing populations but also levying limits to selection ^38^: previous research showed that the two morphs on average disperse similar distances, but the gray morph traveled further in colder winters while the brown dispersed further in warmer winters ^39^. Under increasingly milder conditions, therefore, we would expect phenotypic variation to increase at a higher pace in the brown morph.

We also identified morph-specific differences in the heritable components of plumage coloration as a potential driver of phenotypic divergence. The parent-offspring regression confirmed an overall high heritability (approx. 83%) of plumage coloration in tawny owls ^26,40^. However, genetic determination of coloration appears to be stronger in the gray morph than in the brown morph, as indicated by higher residuals in the linear regression (Fig. 3B) and the significant interaction between parental color scores for gray but not for brown in the additive model (Fig. 5B). Morph-specific heritability estimates Together, both results hint at the existence of morph-specific genetic and potentially non-genetic effects for quantitative color expression. In addition to the observed developmental effects in the recruits, sources of non-genetic variation may include environmental effects in the parents, such as differential parental efforts, where food allocation and reproductive effort may vary according to performance of parental phenotypes ^41,42^, affecting offspring phenotypes differently. Instead, genetic factors would imply that the higher contribution of parental coloration to recruit coloration in the gray morph is due to a stronger genetic determination of coloration. Indeed, plumage coloration of brown recruits was to a significant extent determined by environmental conditions during early-life, which hints at developmental plasticity. Phenotypic plasticity in pigmentation, the ability of a genotype to produce different colored phenotypes in response to the environment, is often observed in invertebrates ^43–45^, but observations from non-model vertebrates and birds are less common, likely due to logistic challenges ^15,46^. Especially in polymorphic systems, the study of morph-specific plasticity is currently confined to butterflies ^47^ and lizards ^16,48^. In tawny owls, plasticity in plumage pigmentation might allow the brown morph to trace optimal phenotypes under a changing environment more quickly ^16,48^, which might be reflected in both the increasing frequency of the brown morph ^26^, but also the relative steadiness of the increase in color scores through the years (Fig. 2A).

Our results support the idea that discrete color polymorphisms are expressed through gene-switches, which are mutations at genes that govern a genetic pathway for the production of a specific pigment in a fraction of the individuals of a population, while inhibiting or largely reducing it in the other fraction ^16,23,49^. While this makes color polymorphisms powerful markers of discrete genetic variation ^2^, gene-switches might not be involved in the mechanisms contributing to continuous intramorph variation, but may posit a physiological limit to phenotypic responses. In some species, melanization in the melanic morphs is strongly correlated with the number of variant protein-coding gene alleles, which seem therefore to act as quantitative trait loci ^23^, whereas, in other systems, different degrees of melanization are associated with physiological processes such as hormonal regulation and dietary concentration of minerals and/or amino acids ^43,50^. In support of this, our parent-offspring regression revealed that approx. 41% of the total phenotypic variance in plumage coloration is due to additive genetic effects (Fig. 3). This suggests that in addition to an overarching Mendelian genetic architecture that determines morph, additional quantitative trait loci and environmental factors may contribute to individual variation in continuous coloration. Current knowledge of the genomic underpinnings of color polymorphism in tawny owls highlights important structural variation in the melanin concentrating hormone receptor gene (MCHR), which codes key physiological responses involved in adaptation to cold ^28^. Thereby, evidence does point towards gradual variation in physiological profiles, especially in the gray morph. In addition, the diverse molecular origin of pheomelanin coloration suggests that its exact genetic architecture may be more complex than phenotypic data alone seems to suggest ^26^. Overall, the role of genetic architectures interacting with the environment during development remains unclear and requires further research, particularly in the context of color polymorphisms and intramorph phenotypic variation.

Despite the importance of genetic color polymorphisms for biological diversity ^51,52^, explicit consideration of intramorph phenotypic variation remains limited in the study of evolutionary processes ^5,12^. Based on our findings, we argue that quantifying intramorph phenotypic variation can increase our understanding of how polymorphisms are maintained ^52,53^, but also how populations persist in space and time ^22,54^. On the one hand, the significant intramorph variation we detected implies that polymorphic species can host larger phenotypic diversity than previously thought, which may increase the evolutionary potential of populations to respond to environmental change ^37,55^. On the other hand, since each morph can become limited in the amount of heritable phenotypic variation, gene-switch-based polymorphisms may also increase the likelihood of producing maladapted forms with reduced adaptive capacity, which might decrease the stability of the polymorphism within the population. Specifically, while the brown morph continues to produce darker phenotypes that have higher fitness in a warming environment, the gray morph appears to be “locked in” at low plumage pigmentation following a series of harsh winters. The expected increased environmental mismatch of these low-pigmented gray individuals in a warming environment might lead to a further reduction of competitive abilities compared to the brown morph, which could eventually displace the gray morph in the population. This highlights the importance of gene flow into the population, which we identified as an important mechanism in the gray morph to produce more pigmented individuals (Fig. S2B). Overall, gauging how these contrasting effects shape the persistence of polymorphic populations remains challenging, and more long-term studies are needed to better understand how continuous variation within color polymorphisms contributes to evolutionary processes.

### Concluding remarks

In summary, our study shows that fitness optima within the two morphs might be contingent on changes in continuous coloration associated with climatic conditions. Importantly, we found support for a higher plasticity of the brown morph, which has likely allowed the spread of the brown morph under climate change in the study population. In contrast, the gray morph might be more constrained by its genetic architecture and be less efficient in tracking environmental changes. While temporal changes in quantitative traits often reflect ecological rather than evolutionary shifts, in polymorphic systems, they can profoundly impact evolutionary dynamics by either preventing morphs from outcompeting each other or accelerating the loss of the less favored morph. This becomes even more important in the context of human-induced environmental change, which may alter tempo and mode of selection on color phenotypes. Our findings thus raise important questions on how color polymorphism might be maintained, among other mechanisms, through changes in the distribution of continuous phenotypes within morphs.

## Methods

### Study area and field protocol

We used data from a population of tawny owls that was collected between 1978 and 2022. The population is monitored using nest boxes and extends over an area of about 500 km^2^, comprising two almost overlapping and equally sized areas in western Uusimaa, Southern Finland (60°15’ N, 24° 15’ E). Part of the area was monitored from 1978 onward and the other one was monitored from 1987 onward (for description of the study area, see ^27,39^). In Finland, tawny owls live in mixed and boreal forests where they promptly breed in nest boxes from late February to late April (median laying date 31 March). In April, all nest boxes were checked to detect breeding attempts and record information on clutch size as well as hatching date. Upon nest inspection, laying date was estimated retrospectively by counting from hatching or by estimating the age of the chicks from their wing length if they had already hatched. During the nestling period, both parents were trapped at the nest box, aged, measured, and ringed (if not) to allow individual identification ^56^. Males and females have similar capture probability ^35,56^. All birds were captured, handled and ringed with an appropriate ringing license.

### Plumage color scoring

The color scores used for studying tawny owls’ color variation were assigned in the field upon capture, following a simple, standardized and repeatable protocol that remained unchanged over the 43 years considered in this study ^35,57^. The scoring protocol focuses on the degree of redness in the plumage, using a continuous semi-ordinal scale ranging from 4 to 14, where a higher score indicates more intense pheomelanic brown pigmentation in feathers, whereas low score indicates gray dominated plumage without any pheomelanic component. The color score is calculated by summing up partial scores made for three different body parts: facial disc (1-3 points), breast (1-2 points), back (1-4 points), and general appearance (1-5 points) ^35^. After scoring, the morphs are assigned based on the overall color scores (4-9 = gray morph, 10-14 = brown morph) ^26,35^. Color scoring effort was constant in time, with only three different observers assigning color scores throughout two sub-populations (Fig. S2). The first sub-population was monitored from 1978-2014 by a single person (observer “a”) who, after thorough teaching and knowledge transfer, passed on all observation-duties (to observer “c”) in 2013. The second sub-population was monitored by a single person from 1987 onward (observer “b”). When owlets fledge it is possible to determine the morph, but not a definitive plumage color score that the individual will have throughout its lifetime because the plumage is not fully developed yet ^26^. Color scores were therefore only assigned to adults upon capture during breeding season, i.e. when individuals were recruited back into the population as breeders. Since tawny owls are long-lived and territorial birds that can occupy the same nest box for many years, a large portion (72%) of all individuals were captured and scored multiple times by the same observer (Fig. S1A). In these cases, color score assignment was performed *de novo*, i.e. observers assigned a score without knowing previous color scores to avoid measurement bias. Using this protocol, color scores were highly repeatable over time ^35^ (see also Fig. S1 in the Supplementary Information), which suggests that intra-individual variation is small and the between-year assessment of the observers reliable.

### Data preprocessing

Prior to all statistical analyses, we filtered unusable observations from the original dataset (Table S1). Specifically, we removed individuals which showed excessive variation in color scores (standard deviation > 1.5; Fig. S1B). Since there are no reports of ontogenetic changes in plumage coloration in tawny owls, we considered within-individual variation as observer-error. Despite the variation stemming from this error was small (Fig. S1C), we decided to remove it entirely by replacing all measured scores with their median. Hence, all presented analyses using color score refer to the per individual median color score. When a color score record of an individual was incomplete, i.e. when a repeatedly scored individual was lacking scores for one or more years, we used the median value of all collected scores to replace all values, including the missing ones. If only a single value was present in an individual’s record, we used this to replace all missing values. Finally, due to the established color scoring protocol allowing for different ranges of possible color scores between the morphs (gray = 6 values, brown = 5 values), we removed all observations with a median color score of 9 (intermediate individuals, *n* = 67) from the dataset so that both morphs had the same number of possible color scores. This procedure also allowed further removal of individuals whose morph assignment might have changed over time. To ensure equally distributed data and homogeneity of variances for statistical analyses across both morphs, we transformed the color scores from an absolute scale to a relative scale ranging from 1 to 5 (Fig. 2A), to which we just refer as “color score” in the main text and in all figures.

### Statistical analyses

We conducted all statistical analyses in the statistical programming language R (v4.2.3 ^58^). To address the first hypothesis (presence of intramorph variation in plumage coloration), we characterized within-morph changes in color score using Generalized Additive Models (GAM; *mgcv* package v1.8-42) ^59^, which are highly suitable to model and test both linear and non-linear patterns in time series. To test the second hypothesis (interactive effects of plumage coloration, winter conditions, and breeding behavior in adults) and the third hypothesis (contribution of developmental effects to color score change) we conducted two sets of analysis. Both approaches used Structural Equation Models (SEM; *lavaan* package v0.6-15)^60^ to test for causal relationships among all relevant variables within a single path diagram, as well as GAMs to test for interactive effects and non-linear relationships in specific paths of the SEMs (linear path analysis cannot test for interactive effects). In addition to the GAMs and SEMs, we conducted a parent-offspring regression using a linear model to estimate heritability of color scores, and visually inspected trends in the residuals as a proxy of transgenerational changes in morph specific coloration and putative non-genetic effects (*stats* package v4.3.3 ^58^). To check whether heritability was temporally stable over the observed period we tested the residuals of the linear regression using an ordinary least squares regression with an autocorrelation term (*nlme* package v3.1-164 ^61^).

#### Variation in intramorph plumage coloration

We implemented a single additive model (GAM 1; Table 1) to test for the presence of morph specific variation, and to visualize the dynamics of color score over time. The GAM included morph as linear term (“parametric coefficient”) and year by morph as non-linear term (“smooth term”), as well as random intercept for each observer-morph combination for the smooths (Fig. S2A). We used two additional GAMs to test for the differences in color score dynamics between residents and dispersing / migrating individuals (GAM S1; Table S2), and for differences between the entire population and the new cohort of each year (GAM S2; Table S2). All GAMs used thin plate regression splines for the smooth terms, 5 knots for the basis function, and restricted maximum likelihood (REML) for the smoothing parameter estimation.

#### Relationship between plumage coloration, breeding behavior and success in adults

We used a path analysis to test for effects of winter conditions on color score in the adult breeding population (*n* = 1972; Table S1), as well as laying date and recruitment success (SEM 1; Table S3). The path diagram includes time (year) and environmental variables between Nov. 1st and April 30th (air temperature [mean, hereafter “temperature”], air temperature variation [coefficient of variation, hereafter “temperature CV”] as a proxy of extreme events, snow depth [mean], snow days [N]), as well as morph, on color score, laying date, and recruitment success in the breeding population. The information on environmental variables was collected by the Finnish Meteorological Institute (FMI) at the weather station near Helsinki-Vantaa airport, situated *ca.* 50 km east of the study area. Laying date in our dataset sets March 31st of each year as the reference (0), and scores deviations from it accordingly (e.g. March 20th = -11, April 3rd = 3). Recruitment success is scored as 1 if any number of fledglings of a breeding pair of adults is recruited back into the population (i.e. appears again in the database), and 0 if not. Morph is coded as a binary variable (0 = gray, 1 = brown), and a positive value would indicate that brown has a stronger positive or negative effect than the gray on the variable it points at. For more comprehensive analysis of three key metrics in adult owls we implemented three additive models, all of which included a morph-specific interaction term: GAM 2 tested for interactive effects of air temperature and snow days on breeding adult color score, GAM 3 tested for interactive interactive effects of air temperature and snow days on laying date, and GAM 4 tested for interactive effects of laying date and color score on recruitment success (see Table 1 for details). All GAMs used thin plate regression splines for the smooth terms, 5 knots for the basis function, and restricted maximum likelihood (REML) for the smoothing parameter estimation.

#### Genetic vs. developmental effects on plumage coloration in recruits

We used a second path analysis to investigate the effects of parental morph and color score, as well environmental dynamics during the breeding period on recruit color score (SEM 2; Table S3). For this analysis we only used the records of recruits with known parental morph, color score and laying date (*n* = 170; Table S1). As a proxy of environmental changes during the rearing phase, we included temperature, temperature CV, snow depth and snow days in a 56 day window after the laying date. This was the average time in our population between laying eggs and the timing of the oldest chick in a brood turning 25 days, shortly prior to fledging. For more comprehensive analysis of parental color scores, as well as environmental conditions on recruit color score, we implemented two additive models, both of which included a recruit-morph-specific interaction term: GAM 5 used an interactive term with maternal and paternal color scores to test differential heritability between morphs and paternal or maternal side. GAM 6 used an interactive term of temperature and temperature to test for morph specific susceptibility for environmental effects on color score development. Both GAMs used thin plate regression splines for the smooth terms, 5 knots for the basis function, and restricted maximum likelihood (REML) for the smoothing parameter estimation.

### Data availability

The data supporting the findings of this study and the codes to reproduce the analyses and the figures will be available in a digital repository upon acceptance.

## Acknowledgements

This paper is dedicated to the memory of Kari Ahola, who passed away during the preparation of the work. Without him, this work would have not been possible. We greatly thank Teuvo Karstinen and the other members of KBP for the many hours spent conducting fieldwork and data sharing. We also thank Glaucia Del-Rio for commenting on a draft of the manuscript. The work was supported by the Swedish Cultural Foundation (grant numbers 168034 and 188919 to P.K.). A.P. was supported by a Margarita Salas fellowship from the University of Seville (MSALAS-2022-22312). M.D.L. was supported by a Marie Sklodowska Curie Individual fellowship awarded by the European Commission (PhenoDim; Grant No. 898932). This is paper number 25 from Kimpari Bird Projects (KBP).

## Author contributions

A.P., M.D.L. and P.K. conceived the study. E.A. and P.K. collected the data in the field. A.P. and M.D.L. analyzed the data. A.P. and M.D.L. wrote the first draft with inputs from P.K.. All authors (A.P., M.D.L., E.A. and P.K.) revised and commented on the manuscript and approved the final version.

## Supplementary Information

### Tables

**Table S1.**
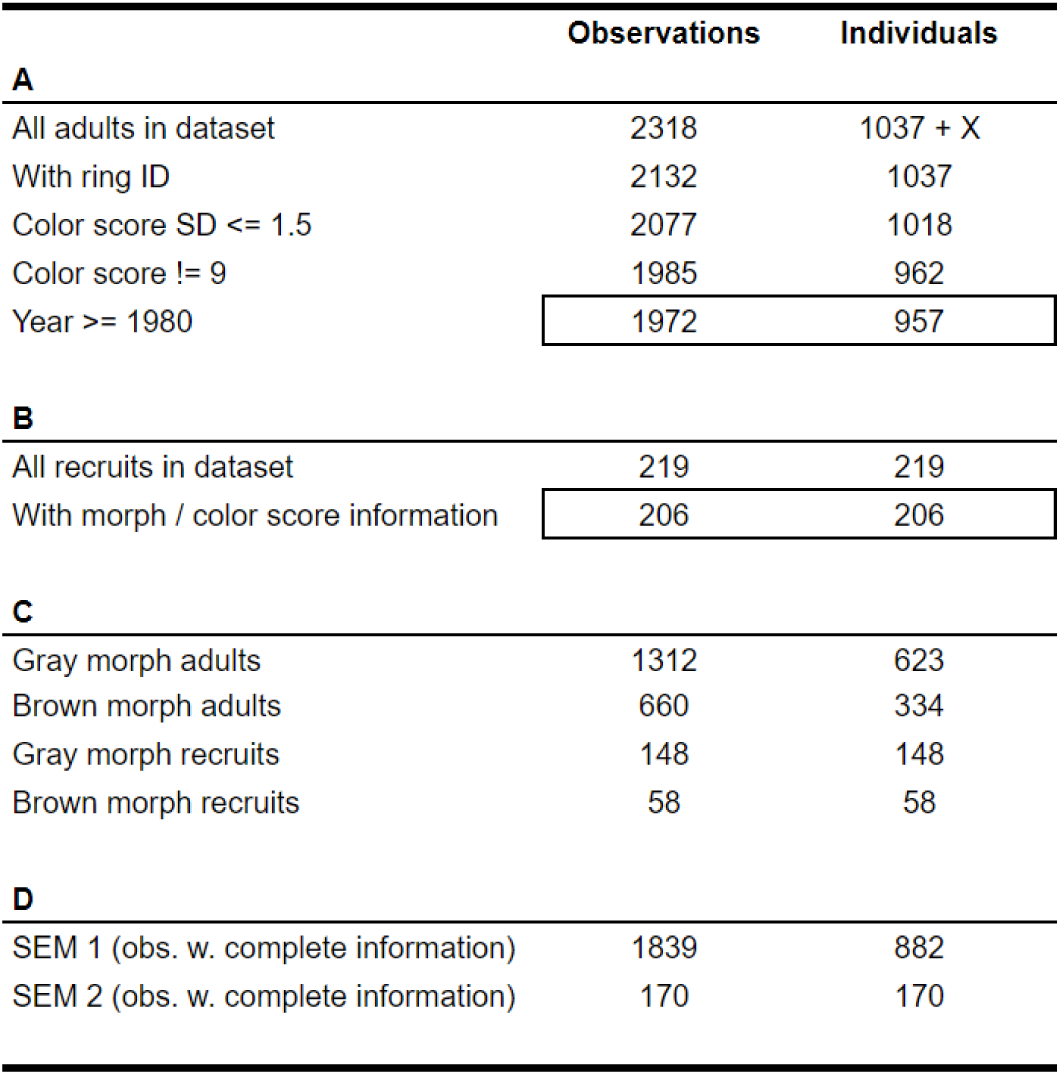
Breakdown of the number of observations. **A:** Adults. Removal of incomplete observations (intersection of all individuals with complete ID records and a standard deviation (SD) for color score that was smaller than 1.5 [see Fig. S1]) left us with 1972 usable records from 957 individuals. **B:** Recruits. Of all recorded recruits, 206 recruits had complete color information. **C:** Number of observations in relation to morph. 68% of the observations (67% of the individuals) in the usable dataset were gray morphs. **D:** Due to missing covariates (e.g. laying date and/or both parent ID) we could not use all observations in the structural equation models (SEMs).

**Table S2.**
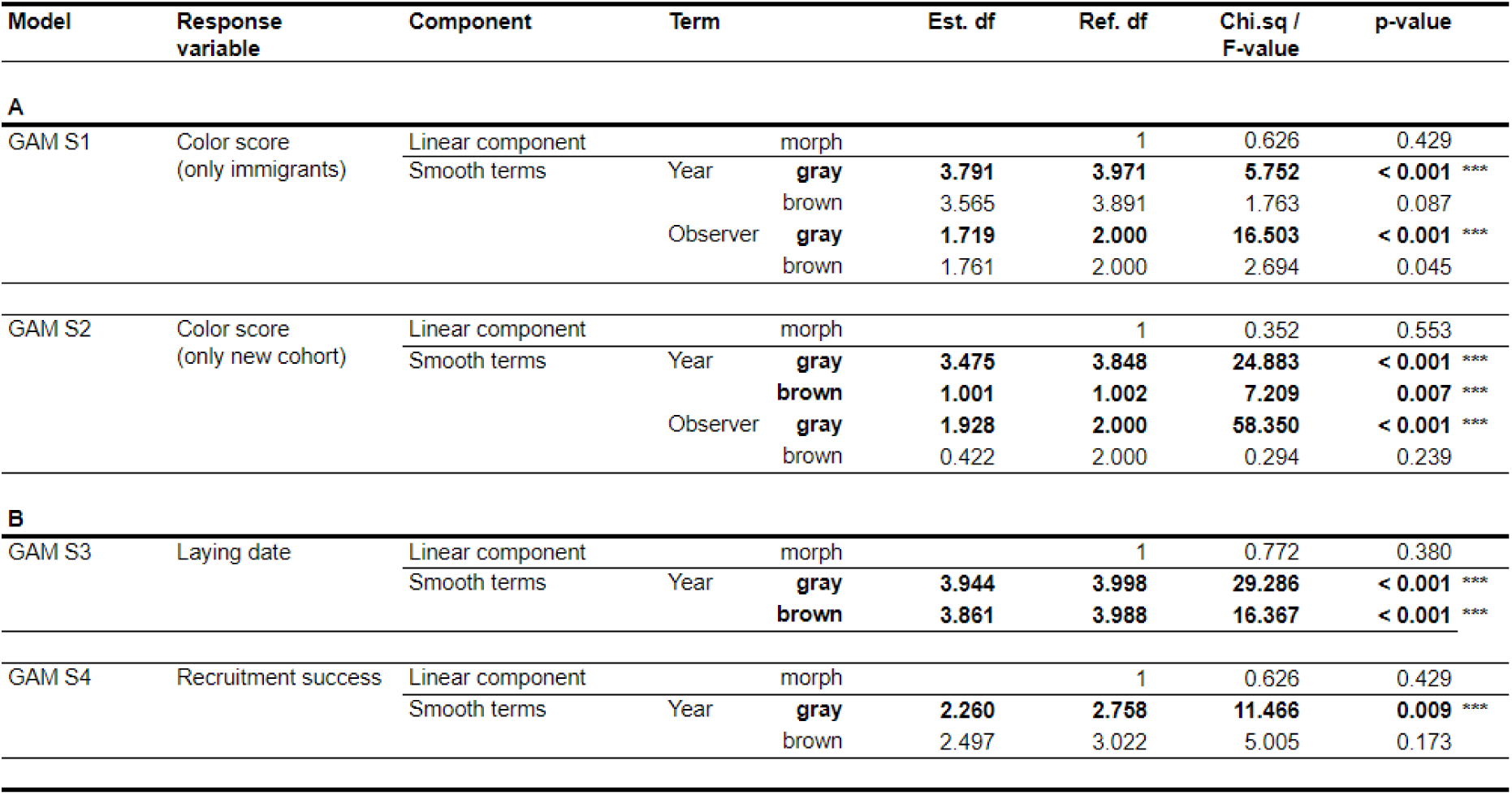
Summary tables for Supplementary Generalized Additive Models (GAMs) **A:** Tests for differences between morphs in color score dynamics of the entire population and immigrants only (GAM S1), as well as the entire population and the new cohort of every year (GAM S2). **B:** Tests for differences between morphs in laying date (GAM S3) and production of recruits (GAM S4) of breeding adults. Significant (0.05 > p ≥ 0.01, *) and highly significant (0.01 > p, ***) results are in bold.

**Table S3.**
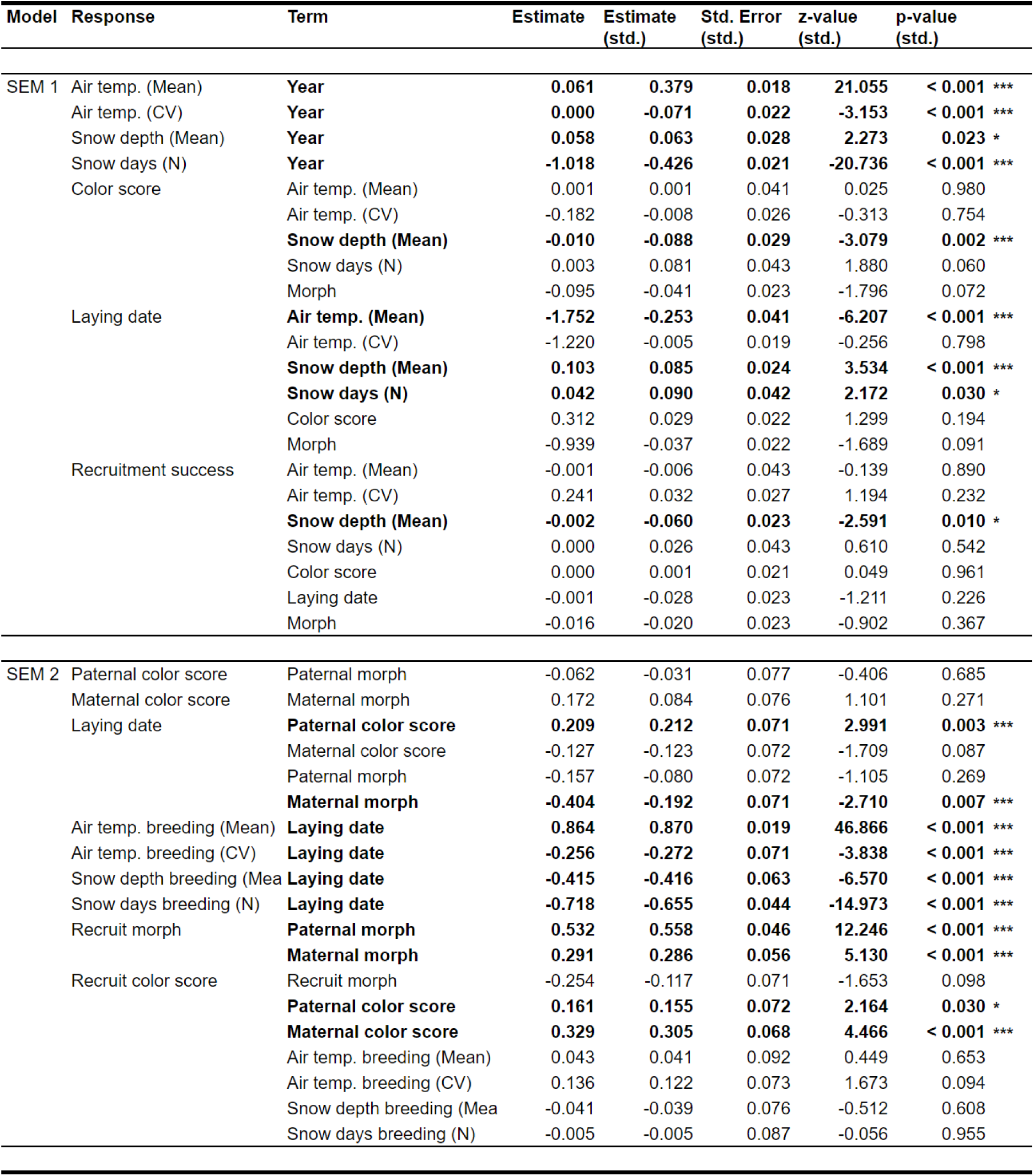
Summary table for the Structural Equation Models (SEM) SEM 1 tested for environmental effects on individual color score, laying date and recruitment success across both morphs (N = 1839). SEM 2 tested for genetic and environmental effects in the recruits (SEM 2, N = 170) through path analysis (see Fig. 4A and 5A). Significant (0.05 > p ≥ 0.01, *) and highly significant (0.01 > p, ***) results are in bold.

### Figures

**Figure S1.**
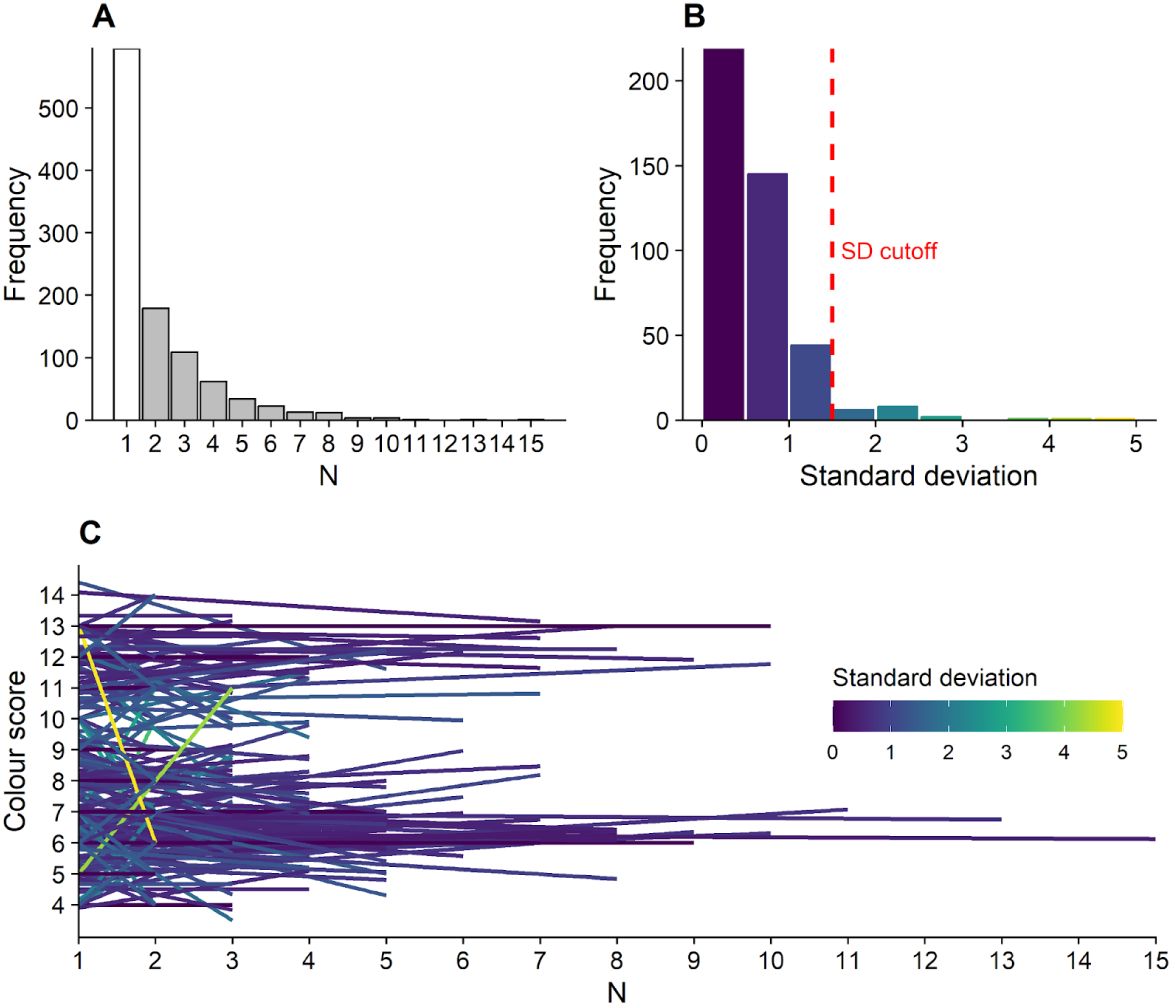
Repeatability of color scoring per individual. **A:** Number of measurements per individual over the years. **B:** distribution of standard deviation (SD) of color scores of individuals that were measured at least two times (gray bars in panel A). Individuals with a standard deviation higher than 1.5 were not used in the analyses (dashed red bar indicates cutoff). **C:** Regression lines of color scores of individuals that were measured multiple times (including the ones that were not used). The color coding denotes the standard deviation across all measurements per individual.

**Figure S2.**
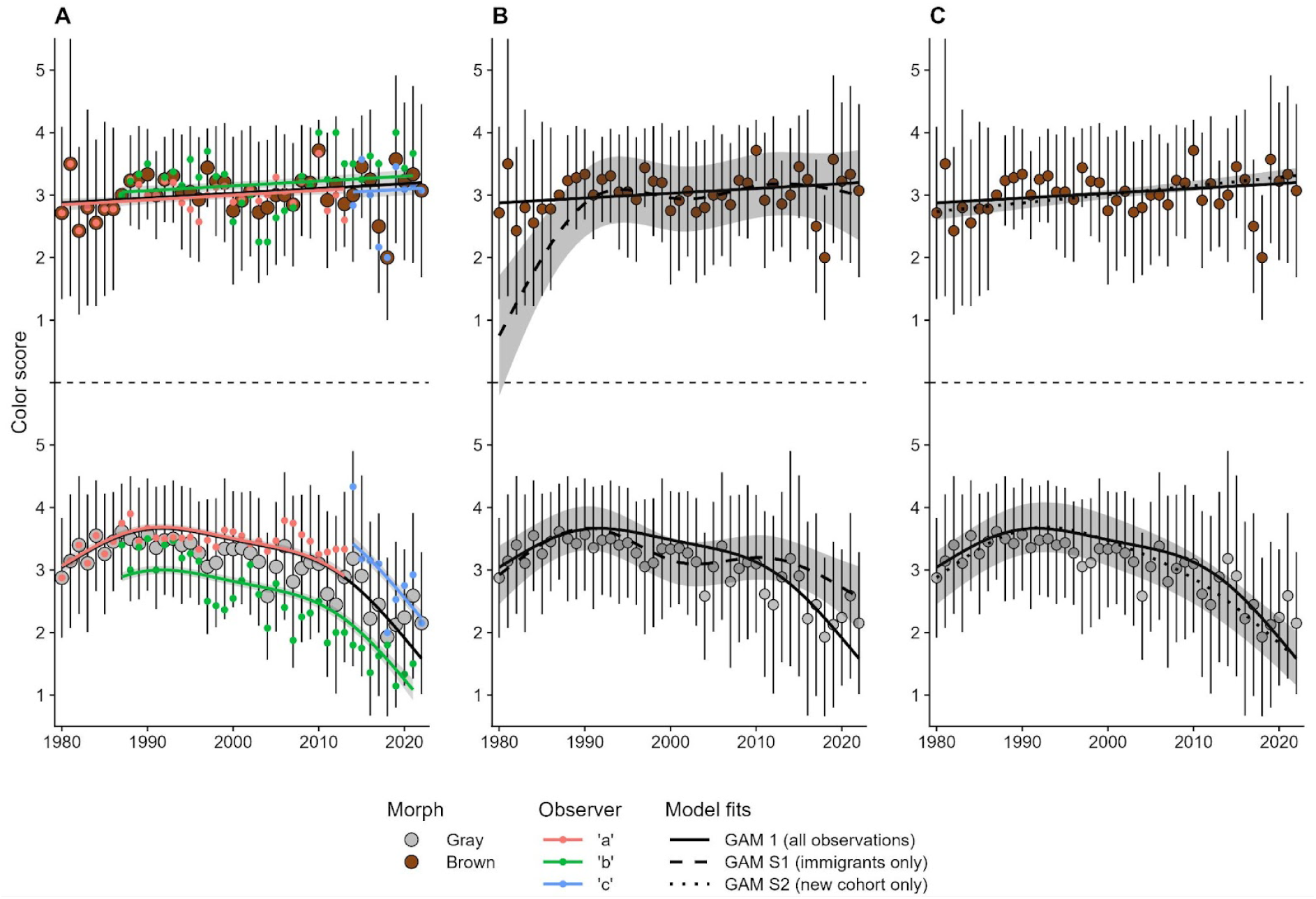
Relative color score dynamics - observer effects and demographic contexts. **A:** Large points denote the yearly average of morph-specific color scores using all individuals per each year (mean ± SD, N = 1972); small three-colored points indicate observer specific averages per year. The solid black line is the model fit from GAM 1 (shading denotes ± SE of fit) with transformed estimates (gray + 3, brown + 9), solid colored lines are fits for observer random intercept effects. **B:** Yearly average (mean ± SD) of morph-specific color scores contrasting the entire population (black solid line) and immigrants (black dashed line; N = 1972). Model fit for immigrants stems from GAM S3; shading denotes ± SE of fit. **C:** Yearly average (mean ± SD) of morph-specific color scores contrasting the entire population (black solid line) and new breeding cohorts for each year (black dotted line; N = 1972). Model fit for new cohorts stems from GAM S4; shading denotes ± SE of fit.

